# Sleep strengthens successor representations of learned sequences

**DOI:** 10.1101/2025.06.11.658893

**Authors:** Xianhui He, Philipp K. Büchel, Simon Faghel-Soubeyrand, Janina Klingspohr, Marcel S. Kehl, Bernhard P. Staresina

## Abstract

Experiences reshape our internal representations of the world. However, the neural and cognitive dynamics of this process are largely unknown. Here, we investigated how sequence learning reorganizes neural representations and how sleep-dependent consolidation contributes to this transformation. Using high-density electroencephalography and multivariate decoding, we found that learning temporal sequences of visual information led to the incorporation of successor representations during a subsequent perceptual task, despite temporal information being task-irrelevant. Importantly, individuals with better sequence memory performance exhibited stronger successor incorporation during the perceptual task. Representational similarity analyses comparing neural patterns with different layers of a deep neural network revealed a learning-induced shift in representational format, from low-level visual features to higher-level abstract properties. Critically, both the strength and transformation of successor representations correlated with the proportion of slow-wave sleep during a post-learning nap. These findings support the idea that sequence learning induces lasting changes in visual representational geometry and that sleep strengthens these changes, providing mechanistic insights into how the brain updates internal models after exposure to environmental regularities.

## Introduction

External experiences continuously shape and update the brain’s internal model of the world. These experiences are dynamic though, and individuals do not merely encode representations of isolated events but also establish structured relationships between them. For example, a child who sees a Welsh Corgi for the first time may encode only a basic representation of a “small, fluffy creature”. However, upon repeatedly observing the Corgi following a girl into a house, the child might begin to associate these elements, gradually constructing a predictive model of their temporal co-occurrence (Fig. 1a). While this example illustrates how we update our world model in daily life, the mechanisms underlying the integration of temporal regularities into existing representations remain poorly understood.

**Figure 1.**
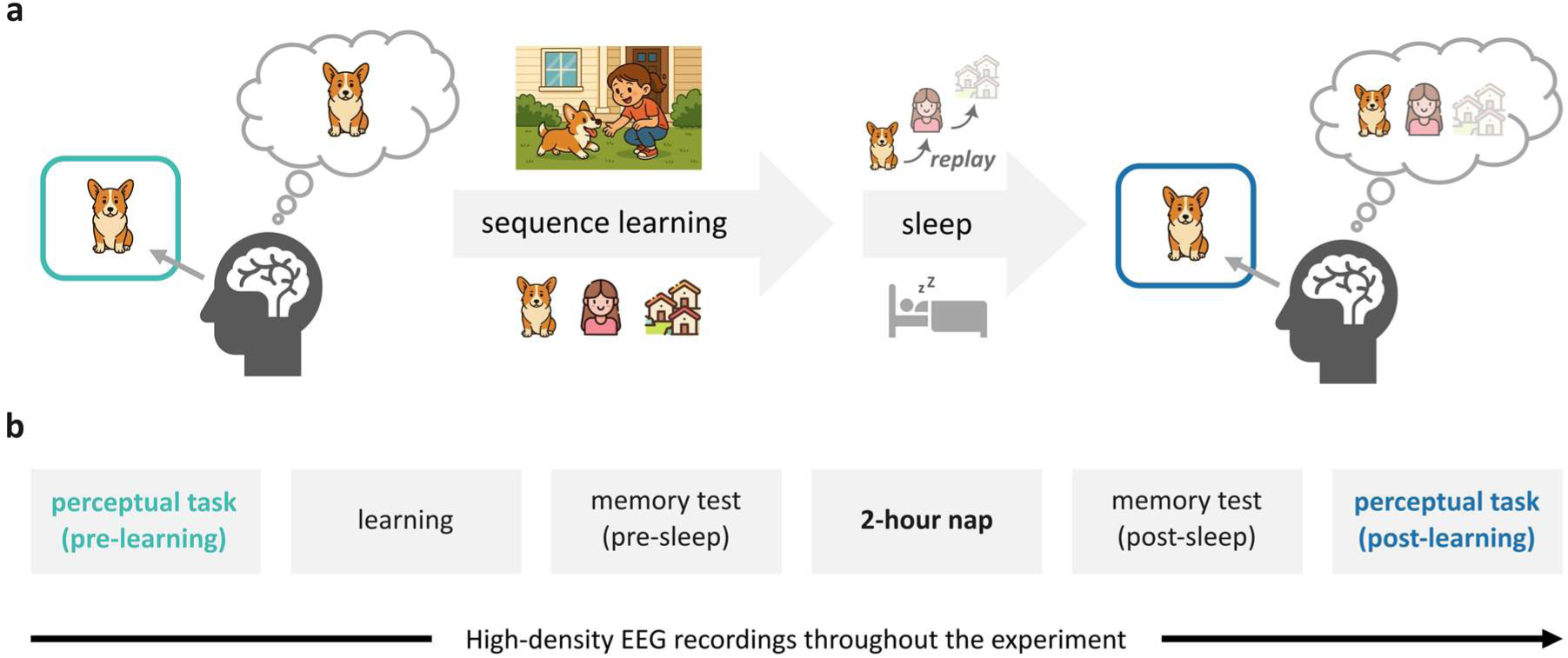
Emergence of Successor Representations and Experimental Design. **(a)** Example of how sequence learning and sleep might change neural representations. Upon encountering a Welsh Corgi, the brain primarily represents the current stimulus entity. If the Corgi is part of a recurring temporal sequence (Corgi → Girl → House), subsequent stimuli (Girl and House) might be integrated into the Corgi representation. Post-learning sleep might provide an opportunity for the brain to replay learned experiences and thereby further strengthen successor representations. Upon post sleep exposure to a Corgi image, brain activation patterns might reflect both the current stimulus (Corgi) as well as learned successors (Girl, House). Faded images indicate weaker representations. Images sourced from AI-generated content and https://www.flaticon.com; **(b)** Timeline of the experiment. Participants first completed a perceptual task, followed by a sequence learning task (Memory Arena). Memory for the learned sequence was then assessed both before and after a period of sleep. Finally, participants completed the perceptual task again.

According to the ‘successor representation’ theory, the brain does not merely represent the current state of the world but also maintains a predictive map, encoding expected future states based on learned temporal transitions (Dayan, 1993; Momennejad et al., 2017; Stachenfeld et al., 2017). This predictive coding mechanism is thought to support flexible behaviour and inference by integrating both immediate states and anticipated future contingencies. While initially developed within the domain of reinforcement learning, recent research suggests that successor representations may also underlie broader cognitive processes, including the organization of human episodic memory (Tacikowski et al., 2024; John et al., 2025; Zhou et al., 2025).

Empirical studies provide supporting evidence that learning temporal regularities across experiences shapes subsequent neural representations. Specifically, representations of temporally proximal stimuli tend to be more similar to one another than to temporally distant stimuli (Greco et al., 2024; Hindy et al., 2016; John et al., 2025; Reddy et al., 2015; Schapiro et al., 2012; Tacikowski et al., 2024). Notably, this effect can emerge after relatively few exposures (Reddy et al., 2015), persists even when such regularities are irrelevant for current task demands (Tacikowski et al., 2024), and correlates with the replay of neuronal activity in the hippocampus during task-free awake breaks (Tacikowski et al., 2024). However, previous studies have not distinguished between representations of past and future information, instead demonstrating that hippocampal activity patterns exhibit greater similarity for temporally adjacent stimuli, regardless of sequence direction (John et al., 2025; Tacikowski et al., 2024). Recent work has further highlighted the bidirectional nature of these representations, showing that both past and future elements of a sequence are encoded (Tarder-Stoll et al., 2024). This raises the question whether sequence learning specifically promotes prospective representations (consistent with the successor representation theory), and whether such changes in neural representations generalise to other contexts where the initial sequence is no longer behaviourally relevant.

The finding that wake replay strengthens successor representations (Tacikowski et al., 2024) moreover begs the question whether sleep, known to constitute a privileged time window for memory reactivation and replay (Buzsáki, 2015; Diekelmann & Born, 2010; Mölle et al., 2004; Niknazar et al., 2022; Wilson & McNaughton, 1994), contributes to the incorporation of successor representations. Interestingly, a recent study has shown that a night of sleep after a real-life sequential learning experience (guided art tour) selectively bolsters temporal order memory while memory for visual-perceptual features steadily declined over the course of up to one year later (Diamond et al., 2025). Importantly, this selective strengthening of temporal memory was predicted by the duration of slow-wave sleep following learning (SWS; Diamond et al., 2025). However, it remains unclear exactly how sleep transforms memory content with regard to perceptual detail vs. higher-level relational structure.

In this study, we investigate whether sequence learning reshapes neural representations such that successor information is incorporated in a subsequent task context where the sequence is no longer behaviourally relevant (Fig. 1b). Specifically, we address three key questions: (1) Does sequence learning reshape stimulus representations to incorporate successor information? (2) What is the representational format of successor representations (e.g., low-level perceptual vs. high-level conceptual)? (3) (How) does post-learning sleep contribute to the incorporation of successor representations? To tackle these questions, we combined behavioural tasks, high-density electroencephalography (EEG) including Polysomnography (PSG) and deep neural network (DNN)-based representational similarity analysis. Our findings revealed that successors of learned images could be reliably decoded in a subsequent non-memory task, with immediate post-learning memory performance predicting the strength of this effect. Additionally, DNN-based representational similarity analysis suggested that successor representations shifted toward high-level visual features after learning. Importantly, greater proportions of SWS predicted the strength of successor representations as well as their shifts towards high-level formats. These results suggest that learning reshapes the representational geometry of visual experiences and that SWS contributes to this transformation.

## Results

Participants first performed a perceptual task, in which 50 unique images from five categories (objects, faces, scenes, letter strings, and body parts; Fig. 2a) were shown in pseudorandom order, with the instruction to press a button when a given image repeated from one presentation to the next (10% of ‘target’ trials, ‘1-back’ task). The perceptual task was followed by the ‘Memory Arena’ task presenting the same images in fixed temporal sequences and spatial locations (Extended Data Fig. 1; Büchel et al., 2024). After reaching a learning criterion, participants took a ~2-hour nap. Memory accuracy was tested before and after sleep, followed by a repetition of the perceptual task. By analysing neural activity during the two perceptual tasks, we examined how successor representations changed with learning. Sleep EEG recordings allowed assessment of how sleep architecture influences these changes.

**Figure 2.**
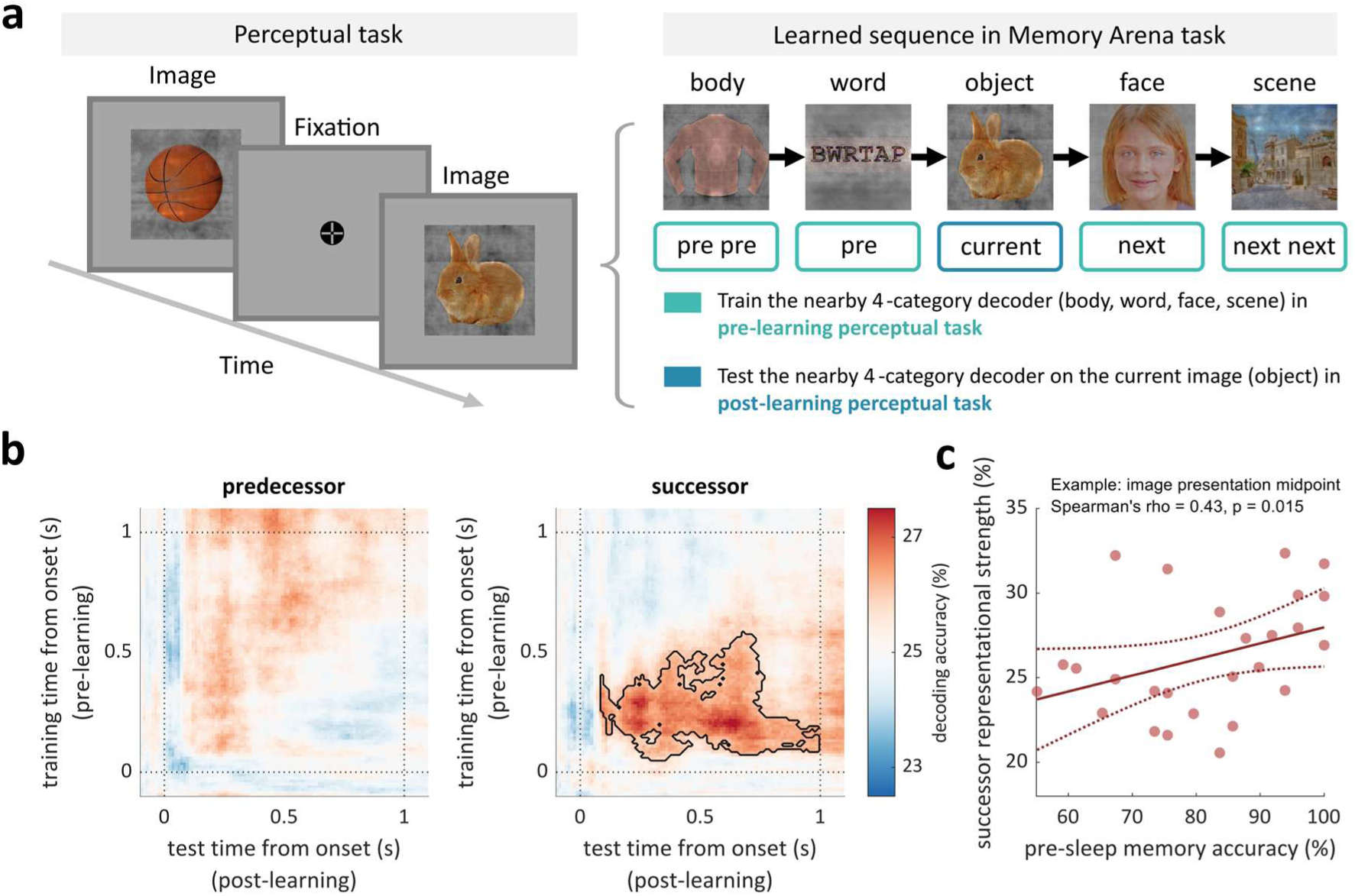
Learning incorporates successor representations. **(a)** Left: example trials from the perceptual task, including image presentation (1000 ms) and fixation period (750-1250 ms). Right: decoding approach for the currently displayed image (bunny). Example image sequence from the Memory Arena task (from previous previous, previous, current, next, to next next around the ‘bunny’ image). Decoders were trained on EEG data from the pre-learning perceptual task using the four nearby categories (body parts, letter strings, faces, and scenes) and were tested on EEG data of the ‘bunny’ (object) image in the post-learning perceptual task. Examples of face stimuli shown were sourced from an AI-generated dataset (https://github.com/RichardErkhov/ai_generated_faces), used under the MIT License; **(b)** Group-level time-by-time decoding accuracies of predecessor and successor images across participants. Black contour indicates significantly higher decoding accuracy than chance level (*p*_cluster_ = 0.028, N = 26, one-tailed cluster-based permutation test); **(c)** Correlation between successor decoding accuracy and immediate post-learning memory accuracy at the *a priori* chosen midpoint of image presentation intervals (0.5s after image onset for both training and test trials). The solid line shows the best-fit linear regression line. The dashed lines indicate the 95% confidence bounds for the fitted line.

### Learning incorporates successor representations

Participants performed well in both the Memory Arena task (mean accuracy ± SD: pre-sleep test: 81.39 ± 13.44%; post-sleep test: 73.15 ± 18.07%; *t*(25) = 3.33, *p* = 0.002) and the perceptual task (mean RT ± SD to target detections: pre-learning: 0.581 ± 0.072s; post-learning: 0.584 ± 0.078s; *t*(25) = −0.52, *p* = 0.604).

To examine whether sequence learning reshapes neural representations of visual experiences, we first assessed how well predecessor and successor information could be decoded from neural activity (Fig. 2a). Specifically, for each image category (e.g., objects), we trained decoders on EEG data from the pre-learning perceptual task using the remaining four categories (body parts, letter strings, faces, and scenes). We then applied these decoders to EEG signals during the post-learning perceptual task to generate a time-by-time matrix of classifier category prediction. This allowed us to assess the evidence for the immediate predecessor or successor image against the chance level (25%). Results revealed that decoding accuracy of successor images was significantly above chance (*p*_cluster_ = 0.028, one-tailed cluster-based permutation test; Fig. 2b). In contrast, the decoding accuracies for predecessor, second predecessor, and second successor images did not exceed chance levels (all *p*_cluster_ > 0.173; Fig. 2b and Extended Data Fig. 2a). Importantly, applying the same decoders to the pre-learning perceptual task revealed no significant effects (*p*_cluster_ > 0.127). In fact, directly comparing successor decoding accuracies between pre- and post-learning perceptual tasks revealed a significant cluster with higher accuracies in the post-learning session (*p*_cluster_ = 0.034, one-tailed cluster-based permutation test; Extended Data Fig. 2b). These results rule out the possibility that the successor representation resulted from perceptual similarities among particular image categories.

Next, we examined whether the strength of successor representation was linked to learning behaviour. We first extracted the successor decoding accuracy at the midpoint of image presentation intervals (0.5s after image onset for both training and test trials, see *Methods*) for each participant and correlated it with their sequence learning accuracy. We found that greater immediate post-learning memory accuracy was associated with greater successor representational strength in the final perceptual task (Spearman’s rho = 0.43, *p* = 0.015; Fig. 2c). Extending this analysis across all combinations of training and test time points, we identified a significant cluster where successor decoding accuracy positively correlated with immediate post-learning memory accuracy (*p*_cluster_ = 0.025; Extended Data Fig. 3a). In contrast, delayed post-sleep memory accuracy showed no significant correlation with successor decoding accuracy (image presentation midpoint: Spearman’s rho = 0.09, *p* = 0.652; cluster-permutation: *p*_cluster_ > 0.281; Extended Data Fig. 3b).

Together, these findings suggest that learning image sequences incorporates successor representations, even when such information is not relevant to the current (perceptual) task. Moreover, better immediate post-learning memory performance predicted stronger successor representations. However, delayed post-sleep memory performance was not significantly associated with successor representations, arguing against active sequence recall during the final perceptual task driving the effect.

### Successor representations shift towards high-level visual information

What is the representational make-up of successor information? To address this question, we applied representational similarity analysis (Kriegeskorte et al., 2008) to compare neural representations with those of a deep neural network (DNN, specifically Alexnet; Krizhevsky et al., 2012). This DNN consisted of seven hidden layers (Fig. 3a), with earlier layers primarily encoding low-level visual features (e.g., colour, contrast), and deeper layers encoding more abstract, higher-level properties (e.g., shape, object identity; Cichy et al., 2016; Goldstein et al., 2022; Güçlü & van Gerven, 2015; Yamins & DiCarlo, 2016). We first constructed an image-by-image EEG similarity matrix at each time point, based on pairwise correlations between neural activity patterns elicited during stimulus processing. This was done for both the pre-learning and post-learning perceptual tasks (see *Methods*; Fig. 3b), and only images remembered in both the pre- and post-sleep memory tests were included in this analysis. These matrices capture the time-resolved representational geometry of visual stimuli in the brain. In parallel, we derived model-based pairwise similarity matrices for successor images based on the learned Memory Arena sequence. This was done separately for each layer of the pretrained DNN, spanning a hierarchy from low- to high-level visual representations.

**Figure 3.**
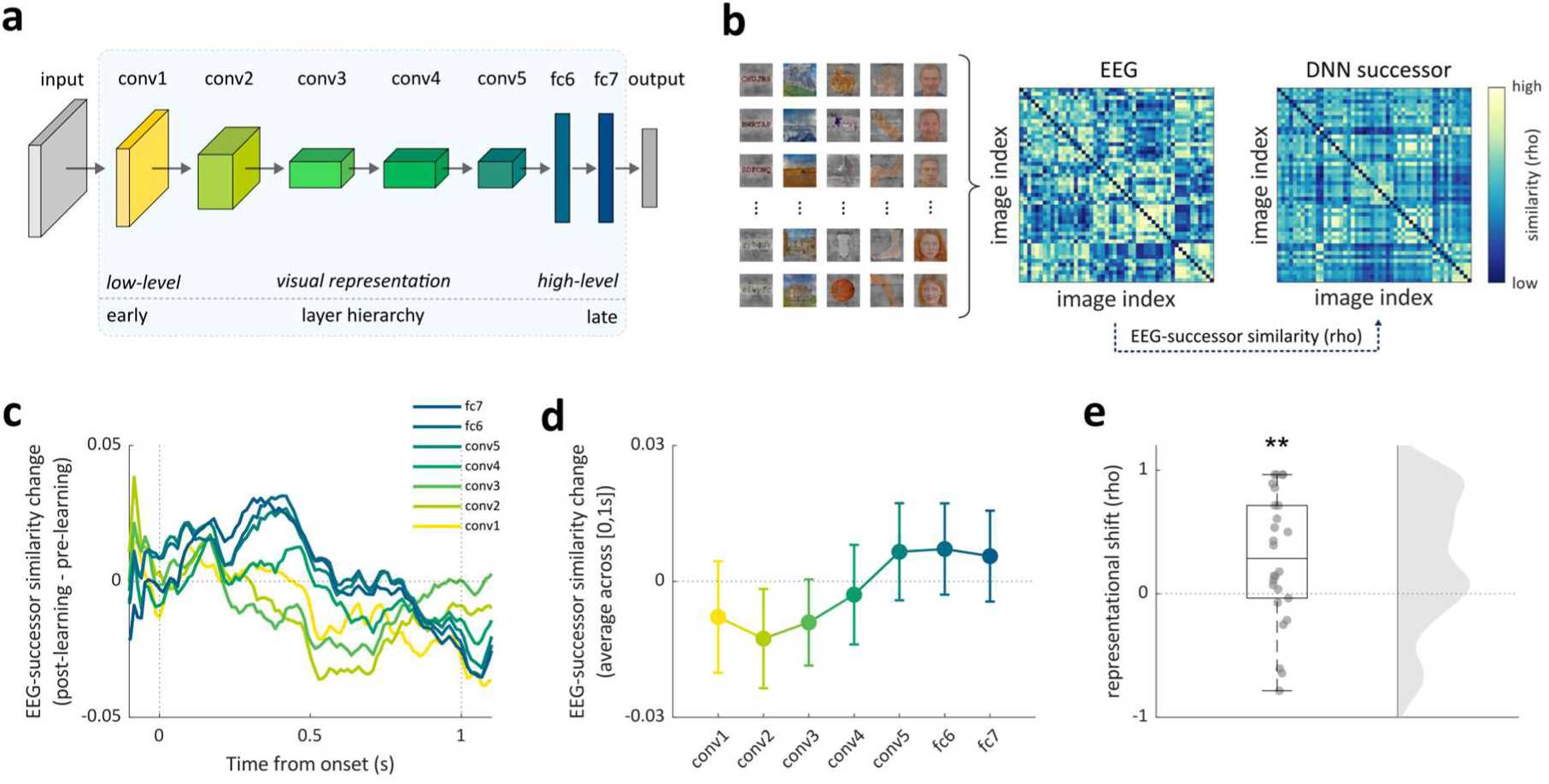
Successor representations shift towards higher-level visual representations. **(a)** Architecture of Alexnet (Krizhevsky et al., 2012). Hidden layers contain conv1-conv5 and fc6-fc7; **(b)** Illustration of EEG-successor similarity analysis. Left: example images used in the Memory Arena task. Middle: Example EEG similarity matrix. Right: Example DNN similarity matrix of successor images in DNN layer 7 (‘fc7’). Examples of face stimuli shown were sourced from an AI-generated dataset (https://github.com/RichardErkhov/ai_generated_faces), used under the MIT License; **(c)** EEG-successor similarity change from pre- to post-learning perceptual tasks across time; **(d)** Group-level EEG-successor similarity change from pre-learning perceptual tasks within the image presentation time window across DNN layers; **(e)** Representational shift: Spearman’s correlations between averaged EEG-successor similarity change and layer hierarchies. Correlations were Fisher z-transformed for statistical testing. ** indicates *p* < 0.01, N =26, two-tailed paired *t*-test.

Confirming the sensitivity of these analyses, we found that both EEG and DNN carried category-specific information, with higher within-category similarity than between-category similarity (Extended Data Fig. 4). To assess the level at which the brain encoded successor information, we correlated neural similarity matrices with DNN-derived similarity matrices at each layer. We hypothesised that neural representations of successor images in the post-learning perceptual task would consist more strongly of higher-level than lower-level DNN features, reflecting a shift toward more abstract representational formats. Consistent with this prediction, comparing EEG and DNN successor similarity matrices in the post-learning perceptual task against the pre-learning perceptual task (as the baseline) revealed a trend toward increased similarity in the deep layers (layer 6: *p*_cluster_ = 0.077; layer 7: *p*_cluster_ = 0.083, two-tailed cluster-based permutation tests; Fig. 3c) and decreased similarity in one of the earlier layers (layer 3: *p*_cluster_ = 0.082).

To quantify this shift towards deep layers, we averaged the change in EEG-successor similarities from pre- to post-learning perceptual tasks within the 1s-image presentation window and correlated them with layer hierarchy (ranging from 1 to 7) for each participant (‘representational shift’ analysis; Fig. 3d and Extended Data Fig. 5a). We found that these correlation values were significantly greater than zero (*t*(25) = 2.88, *p* = 0.007; Fig. 3e). In other words, the change in successor representation format after learning followed the progression from superficial to deep layers in a DNN. A time-resolved analysis also revealed a significant cluster with higher correlation values compared to zero (*p*_cluster_ = 0.018, two-tailed cluster-based permutation tests; Extended Data Fig. 5b).

Lastly, we conducted the same analysis using the current (instead of the successor) image. All DNN layers showed significant correlations with EEG activity (all *p*_cluster_ < 0.001; Extended Data Fig. 6a–b). Interestingly, in both the pre- and post-learning perceptual tasks, we observed that correlation strengths gradually declined from layers conv2 to fc7. To test this observation statistically, we again averaged similarity values across the 1-second image presentation window for each layer and for each perceptual task session. Then, for each participant, we correlated the similarity values of the six deeper layers (conv2–fc7) with the layer hierarchy (ranging from 2 to 7). This revealed a significant negative correlation (*t*(25) = −2.24, *p* = 0.033; Extended Data Fig. 6c), indicating that lower-level visual representations of a DNN were more strongly represented in the EEG data than higher-level representations. This pattern likely reflects the perceptual demands of the task, which required visual discrimination (1-back task, Fig. 2a). However, in contrast to successor images, no significant differences were observed for current images between the pre- and post-learning perceptual tasks (*p*s > 0.357; Extended Data Fig. 6d-e).

In sum, these findings suggest that learning reshapes successor representations and that these representations take on higher-level visual formats, while the sensory encoding of current images emphasises lower-level features and remains largely unaffected by learning.

### SWS predicts successor strength and representational shift

Lastly, we examined how sleep contributes to incorporating successor representations after learning. Nap EEG data were scored with automatic algorithms and manually validated (Fig. 4a; see *Methods*). We then derived the proportion of SWS for each participant (mean ± SD: 24.09 ± 18.07%; Extended Data Fig. 7a) and correlated it with successor representational strength (i.e., decoding accuracy) and shift using both the image presentation midpoint (0.5s after image onset) and in a time-resolved manner. Results showed that higher proportions of SWS were associated with greater successor representational strength across participants (image presentation midpoint: Spearman’s rho = 0.44, *p* = 0.015, Fig. 4b; cluster-permutation: *p*_cluster_ = 0.047, Extended Data Fig. 7d) as well as with the representational shift towards high-level formats (image presentation midpoint: Spearman’s rho = 0.37, *p* = 0.026, Fig. 4c; cluster-permutation: *p*_cluster_ = 0.013, Extended Data Fig. 7e). Importantly, no significant effects were found for any other sleep stage (image presentation midpoint: all *p* > 0.152; cluster-permutation: all *p*_cluster_ > 0.145; Extended Data Fig. 7d-e). Additional analyses ruled out the possibility that these findings were driven by correlations between memory performance and SWS or by interdependence between successor representational strength and shift (Extended Data Fig. 7b-c).

**Figure 4.**
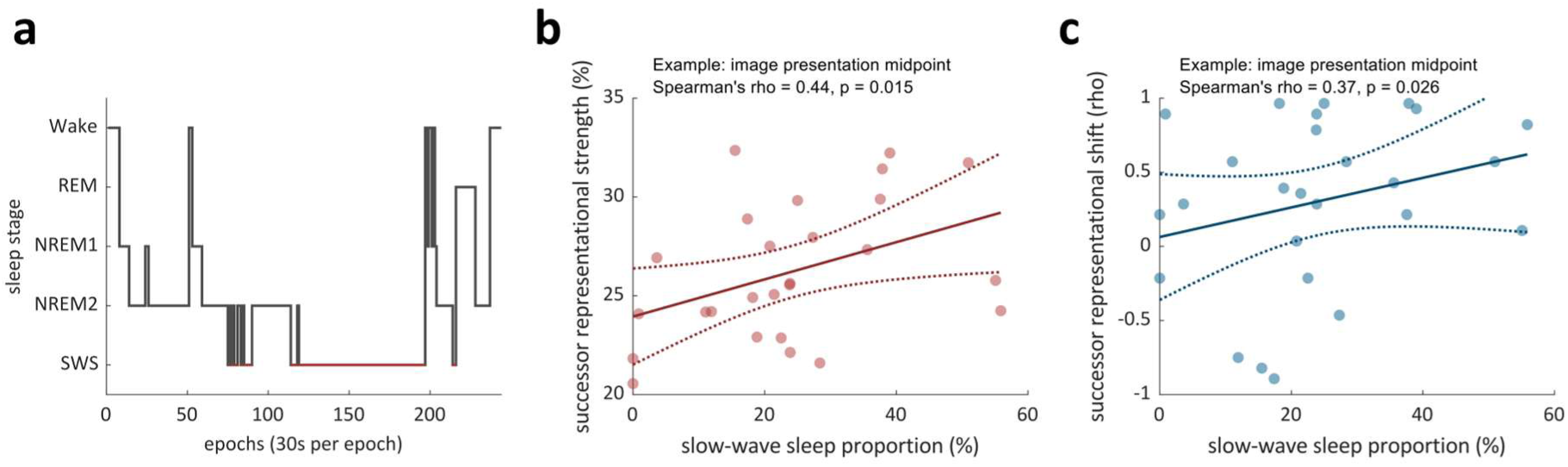
SWS predicts successor representational strength and shift. **(a)** Example sleep hypnogram of one participant. Red lines highlight slow wave sleep (SWS); **(b)** Correlation between SWS proportions and successor representational strength across participants at the image presentation midpoint (0.5s after image onset for both training and test); **(c)** Correlation between SWS proportions and successor representational shift across participants at the image presentation midpoint (0.5s after image onset). The solid line in panels b and c shows the best-fit linear regression line. The dashed lines indicate the 95% confidence bounds for the fitted line.

## Discussion

In this study, we tracked changes in representational geometry before and after learning an image sequence (Fig. 1). By combining high-density EEG recordings and multivariate decoding methods, we found that learning a temporal sequence leads to the incorporation of successor representations in a subsequent non-memory task. Specifically, after learning, the neural activation profiles while viewing an image contained information about its immediate sequence successor, even though sequence information was irrelevant for the current (perceptual) task. Individuals with better pre-sleep memory performance exhibited greater levels of successor integration (Fig. 2). Comparing the representational format of successor information with different layers of a DNN suggested that low-level visual features before learning gave way to higher-level, abstract visual properties after learning (Fig. 3). Crucially, both the strength and shift in format of successor representations were associated with the amount of SWS in a post-learning nap (Fig. 4). Together, these findings suggest that sequence learning incorporates high-level successor representations, and that sleep facilitates this process.

Our findings extend the theoretical framework of successor representations, which were originally proposed in the context of reinforcement learning to integrate current and anticipated future states (Dayan, 1993). More recent work has suggested that this framework may also apply to episodic memory (Gershman, 2012, 2018; Momennejad et al., 2017; Stachenfeld et al., 2017). In line with this view, our results showed robust encoding of successor—but not predecessor—images after sequence learning, even in a task that did not require memory for temporal structure (Fig. 2b). Furthermore, the strength of successor representations correlated with immediate recall performance (after learning and before sleep; Fig. 2c), linking behavioural expressions of sequence learning efficacy to the phenomenon of successor incorporation. The tendency to integrate successive experiences is also consistent with longstanding models of temporal organisation in episodic memory. For instance, the temporal context model (Howard & Kahana, 2002; Polyn et al., 2009) proposes that memory retrieval is guided by a context signal that evolves over time, favouring the recall of temporally subsequent items. Recent computational work has integrated this model with successor representation theory to better capture the predictive structure of memory-guided behaviour (Zhou et al., 2025). Complementing these behavioural and computational accounts, recent human intracranial EEG studies have shown that neurons in the medial temporal lobe encode the temporal structure of visual experiences (Tacikowski et al., 2024; John et al., 2025; Umbach et al., 2020). Although these studies have not distinguished between forward- and backward adjacency, our findings raise the possibility that these temporal codes may primarily reflect forward (successor) information. Future work using intracranial or single-unit recordings could test this hypothesis more directly.

To better understand the format of successor representations, we resorted to tools commonly used in vision neuroscience. Specifically, over the past decade, advances in convolutional DNNs in the domain of computer vision (e.g., Alexnet; Krizhevsky et al., 2012) and natural language processing (e.g., GPT-2; Radford et al., 2019) have provided powerful tools for understanding how the brain organizes complex information. After training, these DNNs form hierarchical representations that progressively transition from low-level sensory features to increasingly abstract, high-level properties—paralleling the structure of human cortical processing (Cichy et al., 2016; Goldstein et al., 2022; Güçlü & van Gerven, 2015; Yamins & DiCarlo, 2016). Using representational similarity analysis (Kriegeskorte et al., 2008), we assessed the alignment of neural activity with different latent layers of Alexnet. Our results revealed a shift in successor representational format: after learning, neural activity patterns became less aligned with low-level visual features and more aligned with high-level, abstract properties (Fig. 3b–d). These findings suggest that learning not only incorporates successor representation but also reorganizes it into a more conceptual format. The observed shift of successor representation complements and extends prior intracranial EEG studies, which revealed a transformation of currently displayed image representations from perceptual to conceptual formats with repeated exposure in short- as well as long-term memory tasks (Liu et al., 2020, 2021; Rau et al., 2025). It further aligns with theories proposing abstraction as a key feature of memory consolidation (Buzsáki & Tingley, 2018; Cowan et al., 2021; Gilboa & Moscovitch, 2021; Kumaran et al., 2016; Nieh et al., 2021; Roüast & Schönauer, 2023).

Intriguingly, we found that the learning-induced changes in neural representations were modulated by sleep. Specifically, the proportion of SWS during a post-learning nap predicted both the strength of successor representations and the extent of their abstraction. These results are consistent with previous work highlighting SWS as a critical window for neural replay, memory reactivation, and representational reorganisation (Buzsáki, 2015; Diekelmann & Born, 2010; Mölle et al., 2004; Niknazar et al., 2022; Wilson & McNaughton, 1994). Although our study did not include a wake control group, recent work offers compelling support for a sleep-specific effect. A study by Diamond et al. (2025) found that participants who slept after real-life sequential learning (a guided art tour) exhibited enhanced temporal order memory but a loss of low-level perceptual detail in behaviour—an effect that was specifically linked to SWS.

This pattern closely parallels our representational-level findings, suggesting that SWS may bias memory consolidation towards abstract, temporally structured information. At the mechanistic level, slow oscillations during SWS are thought to coordinate the timing of sleep spindles and hippocampal ripples (Latchoumane et al., 2017; Staresina, 2024), which in turn supports memory reactivation and consolidation (Schreiner et al., 2021, 2024). This coordinated neural activity may underlie the observed transformation of successor representations. Future studies employing intracranial EEG could directly test whether such cross-frequency coupling facilitates the integration of predictive structures into long-term memory networks.

More broadly, our findings shed light on how the brain updates its internal model of the world. Successor representations provide a mechanism by which experience can reshape predictive structures, allowing individuals to anticipate future events based on learned regularities. This predictive capacity is crucial in everyday contexts where behaviour unfolds over time. An open question in our study, however, concerns the longevity of these representations. Do successor representations persist over time, or do they eventually “wash out” and revert to more veridical representations in the absence of repeated/continued learning? Investigating the durability and stability of these effects over longer timescales remains an important direction for future work.

In conclusion, our findings show that sequential learning induces successor representations in the human brain, even in a perceptual task unrelated to sequential information. These representations shift from perceptual to abstract formats and are supported by post-learning sleep, particularly slow-wave sleep. Together, these results advance our understanding of how the human brain integrates temporal structure into our internal world model.

## Methods

### Participants

Thirty healthy adults (9 males; mean age = 25 years; range = 19–39 years) participated in the study. They received either course credit or monetary compensation for their participation. To ensure sleep-impairing factors did not confound the results, participants were screened for the following exclusion criteria: engagement in night shift work within the past year, recent travel across time zones (within the past two weeks), current use of medications affecting sleep, any history of neurological, psychiatric, or sleep disorders and regular consumption of more than one cigarette per day. Sample outliers were detected using MATLAB’s *isoutlier()* function applied to performance during the final learning block and pre-sleep accuracy scores. This resulted exclusion of 4 participants and a final sample of 26 participants included in the reported analyses. All participants provided written informed consent, and the study was approved by the University of Oxford’s ethics committee (approval code: R85832/RE001).

### Experimental design

Participants first performed a perceptual task including 50 unique images, followed by the ‘Memory Arena’ task presenting the same images in fixed temporal and spatial sequences (Büchel et al., 2024; for an earlier version of the task, see also Petzka et al., 2021). After reaching a learning criterion, participants took a ~2-hour nap. Memory accuracy was tested before and after sleep, followed by a repetition of the perceptual task. EEG recordings were applied throughout the experiment.

The perceptual task included 50 images from five categories (objects, faces, scenes, letter strings, and body parts). Specifically, participants performed a one-back repetition detection task, pressing the ‘down arrow’ key whenever an image was repeated. Each session comprised 660 trials (including 10% repeated target trials), and participants performed the task with high accuracy (mean ± SD: 98.32 ± 0.74%). For subsequent analyses, only correct trials involving non-repeated images were included. Importantly, the image presentation order in the two perceptual task sessions was pseudo-randomised and unrelated to the sequences learned in the Memory Arena task, enabling us to isolate the effects of sequence learning on visual brain representations.

In the Memory Arena task, participants learned the sequential and spatial structure of 50 images across repeated learning cycles (Extended Data Fig. 1, detailed in Büchel et al., 2024). These 50 images were organized into 10 subsequences of five images each, following one of two fixed category orders: (i) letter string, scene, object, face, or (ii) object, scene, letter string, face, with body part images randomly inserted to obscure the primary category sequences. The two subsequence types were counterbalanced across participants. Each learning cycle consisted of two exposure blocks, where images were presented sequentially at their respective locations, followed by a test block in which participants reconstructed the sequence and spatial layout. Learning continued until participants reached at least 66% accuracy in selecting correctly ordered image pairs during a test block or a maximum of 60 minutes had elapsed.

Following the learning phase, participants performed a 5-minute attention task. They were instructed to fixate on a central cross and count each instance it turned dark grey, while ignoring instances where it turned light grey. After that, participants completed the same memory test as in the Memory Arena task, which served as an assessment of pre-sleep memory performance. They were then given a 2-hour nap opportunity (mean sleep duration ± SD: 71 ± 28 minutes), during which polysomnographic data were recorded to monitor sleep stages. Upon waking, participants engaged in an interference task. In this phase, they re-encoded the same set of 50 images, but with both the sequence and spatial locations altered. Each image’s new location differed from its original location by at least 5 pixels (Euclidean distance). The encoding procedure for this new configuration mirrored that of the initial learning phase, but consisted of only one round. Following the interference encoding, participants were asked to recall the new sequence and spatial arrangement. Next, they completed a memory test to retrieve the originally learned sequence and layout, which served as an assessment of post-sleep memory performance. This post-sleep assessment was followed by a final repetition of the perceptual task.

### EEG recording and preprocessing

EEG data were recorded using a 64-channel Brain Products system at a sampling rate of 500 Hz. Electrodes were positioned according to the international 10–20 system, with FCz as the recording reference and AFz as the ground. Six electrodes were reassigned for auxiliary recordings: two for mastoids, two for electromyography (EMG), and two for electrooculography (EOG). This configuration left 58 channels available for scalp EEG recording. Our analyses focused on the EEG data collected during the perceptual task sessions.

Preprocessing was conducted using the following procedures: First, data were downsampled to 250 Hz and high-pass filtered at 0.1 Hz. Line noise in the 49–51 Hz range was removed, and data were re-referenced to the common average. Channels identified as noisy during visual inspection were interpolated; in total, 11 bad channels were replaced across 26 participants.

The filtered EEG signal was then segmented into epochs ranging from 500 ms before to 1500 ms after image onset. Independent component analysis (ICA) was performed on these epochs to identify and remove components reflecting eye blinks, based on visual inspection. The cleaned data were smoothed using a 200 ms moving mean window (MATLAB’s *smoothdata* function), baseline corrected using the 200 ms pre-stimulus interval, and finally z-scored across trials using MATLAB’s *normalize* function.

### Successor representation decoding

To investigate whether sequence learning changes visual brain representations, we conducted multivariate pattern classification analyses using EEG voltage signals from all channels. Within each participant, we first averaged the EEG signal in sliding time windows of 20 ms (five data points) with a step size of 12 ms (three data points). The resulting voltage patterns across all channels served as input features for a multiclass linear discriminant analysis (LDA), implemented using the MVPA-Light toolbox (Treder, 2020).

To assess whether learning induced representations that incorporated sequential structure— such as successors—we examined how well neural activity during post-learning perceptual task trials reflected category information from nearby positions in the learned Memory Arena sequence. For each category (e.g., objects), we trained the classifier using EEG data from the other four categories (e.g., body parts, letter strings, faces, and scenes) in the pre-learning perceptual task session and tested it on untrained object images in the post-learning perceptual task session. This cross-session generalisation design allowed us to identify changes in neural representations that emerged specifically as a result of learning, while minimizing concerns of overfitting or confounding temporal proximity effects. We then compared the predicted category label with the category of neighbouring items in the sequence participants had learned during the Memory Arena task. For instance, if participants had learned a subsequence such as “letter string → object → face,” we examined whether EEG patterns during object trials were more likely to be classified as the successor category (face) or the predecessor category (letter string). This procedure yielded an accuracy estimate for each time point that the decoded category matched the immediate predecessor or successor in the learned sequence. These accuracies were averaged across trials to generate a time-by-time matrix of classifier predictions, which were then compared against the chance level (25%) using cluster-based permutation statistical tests (see Statistical Analysis). We also extended this analysis to more distal sequence positions—examining decoding accuracies for the second predecessor and successor—to further probe the spread of sequence-related representations (Extended Data Fig. 2a).

To relate successor representation to memory performance, we correlated decoding accuracies with pre- or post-sleep sequential memory accuracies across participants at each training and test time point.

Lastly, to confirm that particular effects emerged as a consequence of learning, we applied the same decoding procedures to pre-learning perceptual task data. Successor decoding accuracies from the pre-learning perceptual task session were then compared to those from the post-learning perceptual task session using the same statistical approach. The absence of sequence-based classification patterns prior to learning, along with a significant increase in successor decoding after learning, served as evidence that the observed representational changes were specifically induced by sequence learning occurring in between (Extended Data Fig. 2c).

### Representational similarity analysis

To investigate the format of the successor information, we compared neural representations with those derived from a DNN (Alexnet; Krizhevsky et al., 2012). Alexnet is pretrained on the ImageNet dataset to perform object classification, and has been shown to exhibit a hierarchical organisation of visual representations similar to the ventral visual stream in humans and other primates (Cichy et al., 2016; Goldstein et al., 2022; Güçlü & van Gerven, 2015; Yamins & DiCarlo, 2016). Specifically, earlier layers (e.g., conv1–conv3) predominantly encode low-level features such as edges and textures, whereas deeper layers (e.g., fc6–fc7) capture increasingly abstract and categorical information (Yamins & DiCarlo, 2016).

For each image used in the experiment, we extracted activation patterns (expressed as feature vectors) from each layer of Alexnet. These patterns were used to construct representational similarity matrices by computing pairwise Spearman’s correlations between feature vectors across images. Two types of DNN similarity matrices were created:

1. DNN current similarity matrices: computed using the features of the currently displayed image.
2. DNN successor similarity matrices: computed using the features of each image’s successor (i.e., the image that followed in the learned sequence), rather than the image itself.

After deriving DNN similarity matrices, we next computed EEG-based similarity matrices. For each participant, we selected images whose successors were correctly identified in both memory tests, ensuring reliable recall. Using these remembered images, we computed pairwise Spearman’s correlations of EEG voltage patterns across channels, using the same sliding time window approach as in our decoding analyses.

To validate these similarity matrices, we assessed category-level fidelity by comparing within- and between-category similarities. Both DNN and EEG matrices showed significantly higher within-category similarity, confirming that category-level structure was present (Extended Data Fig. 4).

We next assessed the correspondence between EEG and DNN representations using the same subset of images for each participant’s analyses. First, we examined the correlation between EEG and the DNN current similarity matrix for each layer (EEG–current similarity). This analysis replicated prior findings showing a strong alignment between EEG activity during perception and DNN-derived representations (Extended Data Fig. 6a–b; Liu et al., 2020, 2021; Rau et al., 2025). To evaluate whether EEG activity differentially reflected low- or high-level DNN features, we first averaged, for each participant, EEG–current similarity across the two perceptual task sessions and across the 1-second image presentation window. Then the averaged layer-specific EEG–current similarity was correlated with DNN layer hierarchy (ranks of conv2 to fc7). Positive values indicated stronger alignment with higher-level layers, and negative values indicated alignment with lower-level layers. Correlations were Fisher z-transformed and correlation values equal to ±1 were winsorized to ±0.99 to avoid infinite values during Fisher z-transformation.

To examine how learning influenced neural representations, we assessed the similarity between EEG and DNN successor representations (EEG-successor similarity). To minimize the influence of current image representations in the EEG data, we regressed out the DNN current matrices from the EEG similarity matrices and correlated the residuals with the DNN successor similarity matrices. This correspondence was compared before and after learning for each DNN layer, using cluster-based permutation tests to correct for multiple comparisons across time. Additionally, to quantify learning-related changes during stimulus presentation, we averaged the similarity differences across the 0–1 s post-onset window.

Finally, to quantify representational shifts across the DNN hierarchy (conv1 to fc7), we correlated the magnitude of EEG–successor similarity changes with layer hierarchy (ranks 1 to 7). This was done both across the entire time window and for the average within that window. As before, correlations were Fisher z-transformed and winsorized to ±0.99.

### Sleep architecture and its relation to successor representations

Sleep staging for each participant was performed based on 30-second epochs, using EEG polysomnographic recordings in accordance with the American Academy of Sleep Medicine (AASM) guidelines. Stages included wakefulness, NREM1, NREM2, NREM3 (SWS), and REM sleep. Initial automated staging was conducted using two AI-based tools, YASA (Vallat & Walker, 2021) and Somnobot (https://somnobot.fh-aachen.de). The transitions between stages and any discrepancies between the two automated outputs were then carefully reviewed by two experienced sleep researchers, who finalized the classification of each epoch.

Following staging, we calculated the proportion of time each participant spent in each sleep stage. To examine the relationship between sleep architecture and successor representations, we correlated the proportion of each stage with (i) the successor representational strength (decoding accuracies) obtained from the decoding analysis and (ii) the successor representational shift obtained from the representational similarity analysis.

### Statistical analysis

To assess the temporal profile of successor representations, we used cluster-based permutation testing (implemented in FieldTrip; Oostenveld et al., 2011) to correct for multiple comparisons across time in both decoding and representational similarity analyses. This nonparametric approach identifies clusters of consecutive time points that exceed a predefined threshold (here: *p* < .05), with cluster-level significance evaluated through 2,000 random permutations of the data. For decoding analyses, we used one-tailed tests, as only above-chance effects were theoretically meaningful. For representational similarity analyses, we used two-tailed tests to allow for both positive and negative effects.

To examine the relationship between representational changes and behavioural performance or sleep architecture, we computed correlations between the magnitude of representational change and either sequential memory accuracy (pre- and post-sleep) or the proportion of time spent in each sleep stage, across participants. Correlations were computed at each time point, and the resulting statistical maps were corrected for multiple comparisons using the same cluster-based permutation procedure. We used one-tailed testing for these analyses, based on the hypothesis that better memory performance or a greater proportion of time in slow-wave sleep would be associated with stronger neural representations.

For clearer visualisation and summary of the correlation analyses, we additionally focused on an a priori defined midpoint of the image presentation interval—500 ms after image onset. This time point has previously been associated with both perceptual and conceptual processing (Liu et al., 2021) and thus represented an unbiased choice for evaluating perceptual/conceptual features. At this time point, we evaluated correlation significance using a permutation procedure designed to account for variability in small samples. Specifically, we generated a null distribution by randomly shuffling the correspondence between the two variables across participants 2,000 times and determined p-values based on the proportion of permutations in which the permuted correlation exceeded the observed value.

## Data and code availability

Data and code for this project will be available upon publication on the Open Science Framework (https://osf.io/utdgv/).

## Acknowledgements

This project was funded by the European Research Council (ERC) under the European Union’s Horizon 2020 (grant agreement no. 101001121) awarded to B.P.S and was supported by Medical Sciences Graduate School Studentship at the University of Oxford awarded to X.H. as well as funding support from The Royal Society (project NIF\R1\221006).

## Author contributions

Conceptualisation, X.H. and B.P.S.; methodology, X.H., S.F.S., M.K., P.K.B., J.K., and B.P.S.; validation, X.H.; formal analysis, X.H.; investigation, X.H.; writing – original draft, X.H.; writing – review & editing, X.H., S.F.S., M.K., P.K.B., and B.P.S.; supervision, funding acquisition, B.P.S.

## Extended data

**Extended Data Figure 1.**
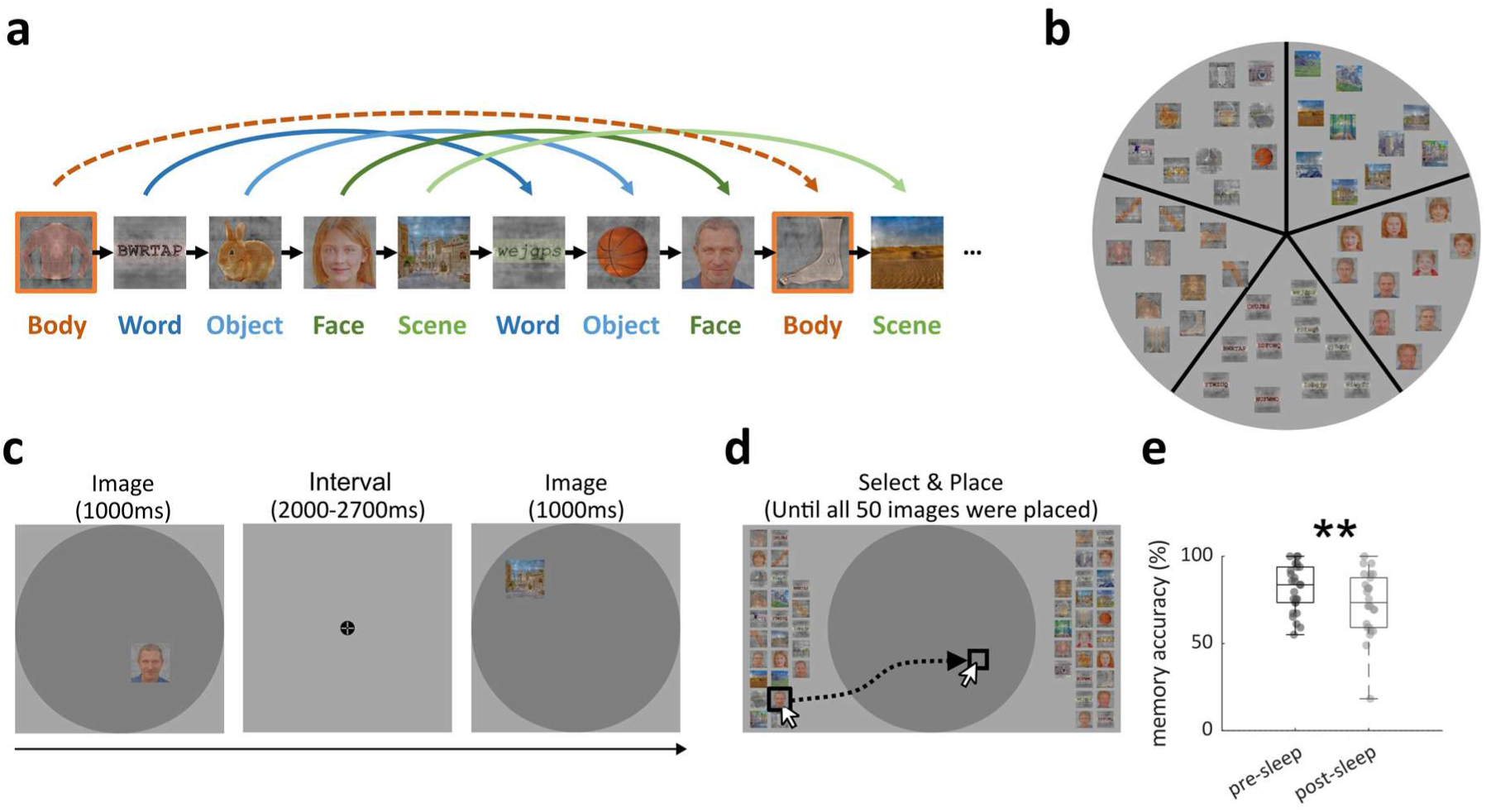
Task and behavioural results. **(a)** Memory Arena sequence design. Participants (N = 26) were tasked with learning the spatiotemporal structure of 50 images. These images belonged to five distinct categories (letter strings, scenes, objects, faces, and body parts) and were organized into 10 subsequences of five images each, following one of two fixed category orders: (i) letter string, scene, object, face, or (ii) object, scene, letter string, face, with body part images randomly inserted to obscure the primary category sequences. The two subsequence types were counterbalanced across participants; **(b)** Memory Arena location design. The Arena was spatially organized into five principal ‘slices’, with each slice corresponding to one of the five main image categories. Each main slice was subdivided into two sub-slices, corresponding to the two subcategories of the main category; **(c)** Exemplary learning trial. Images were presented sequentially at their unique positions. Participants were asked to learn the image and its location; **(d)** Exemplary test trial. All 50 images were randomly distributed around the Arena. Participants selected each image by clicking on it and then click exact position within the arena where they remembered the image had been located. Examples of face stimuli shown in panels a-d were sourced from an AI-generated dataset (https://github.com/RichardErkhov/ai_generated_faces), used under the MIT License; **(e)** Memory accuracy in pre- and post-sleep test. Each dot represents one participant. ** indicates significant difference in memory accuracy between pre- and post-sleep test (*p* = 0.002, N = 26, two-tailed paired *t*-test).

**Extended Data Figure 2.**
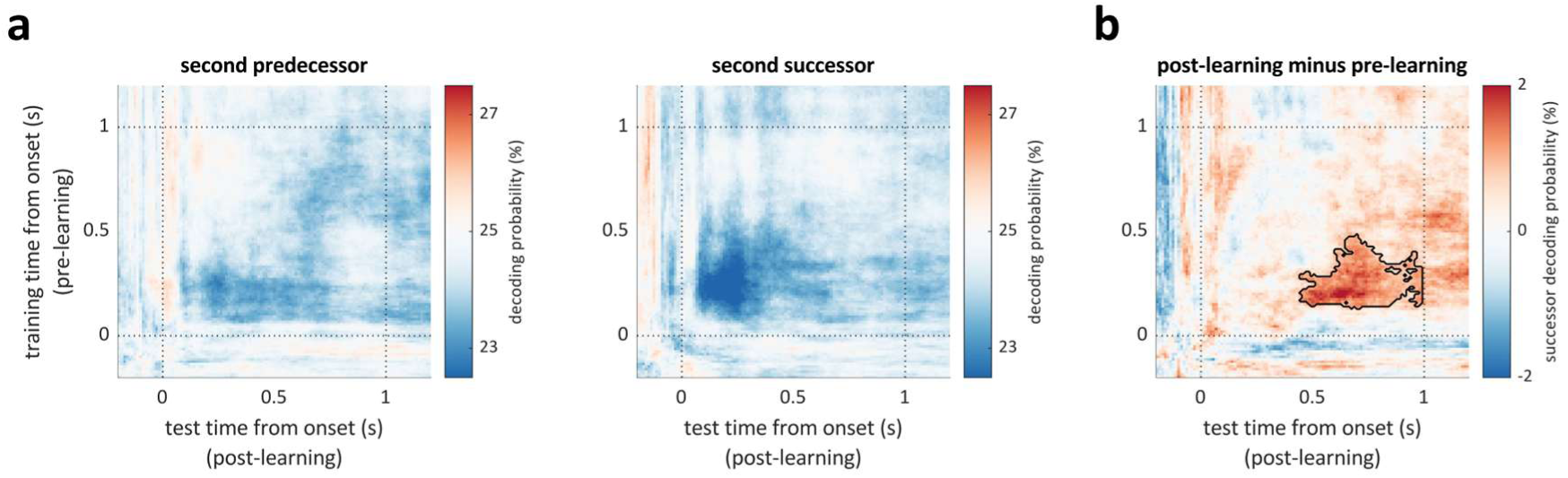
Decoding results of second predecessor and successor. **(a)** Group-level time-by-time decoding accuracies of second predecessor (left) and second successor (right) images across participants (N = 26). Decoders were trained in each time point in the pre-learning perceptual task and were then tested in the post-learning perceptual task; **(b)** Comparison of successor decoding accuracies between the pre- and post-learning sessions. Black contour indicates significantly higher correlations than zero (*p*_cluster_ = 0.034, N = 26, one-tailed cluster permutation tests).

**Extended Data Figure 3.**
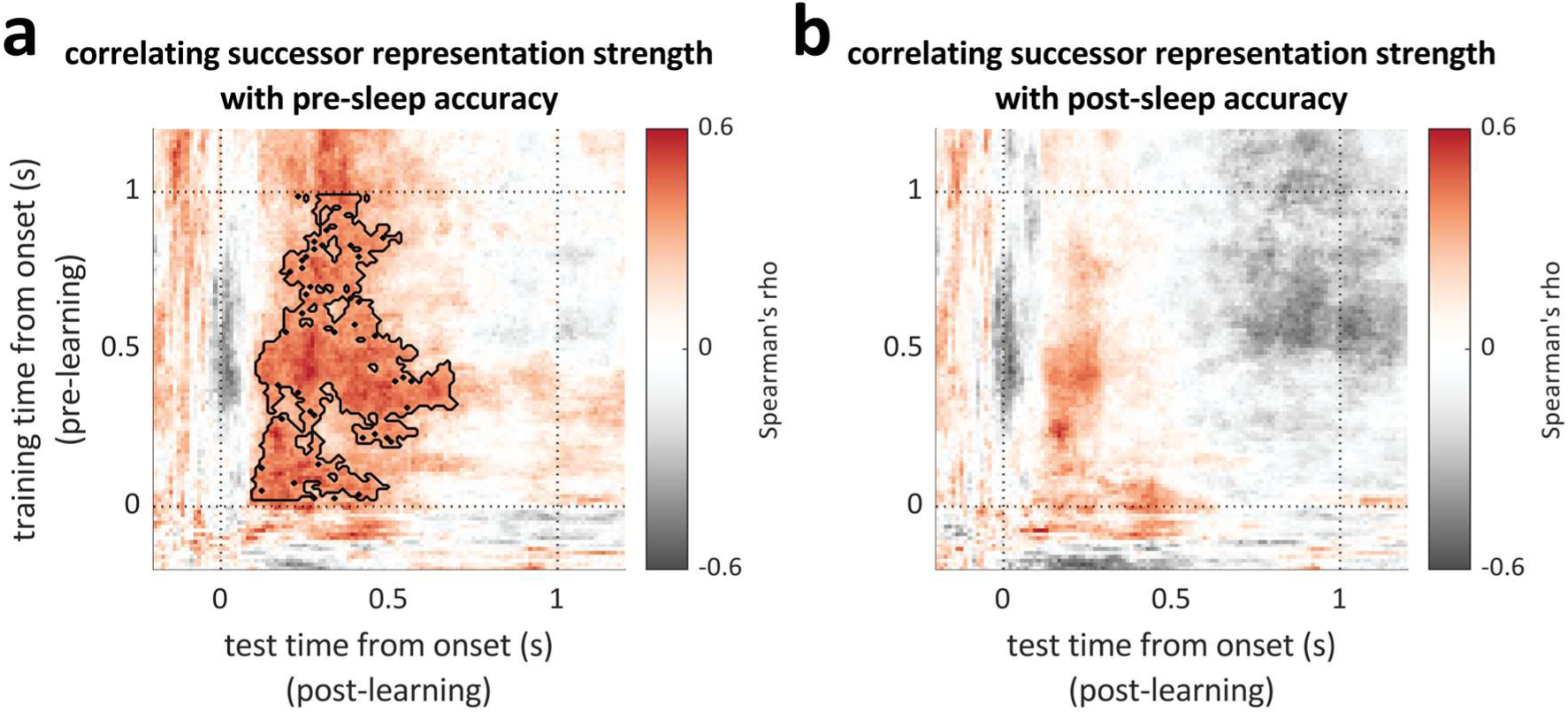
Correlation between successor representational strength and behaviour. **(a)** Correlation between successor decoding accuracy and pre-sleep memory accuracy across participants. Black contour indicates significantly higher correlations than zero (*p*_cluster_ = 0.025, N = 26, one-tailed cluster permutation tests); **(b)** Correlation between successor decoding accuracy and post-sleep memory accuracy (*p*_cluster_ > 0.281, N = 26).

**Extended Data Figure 4.**
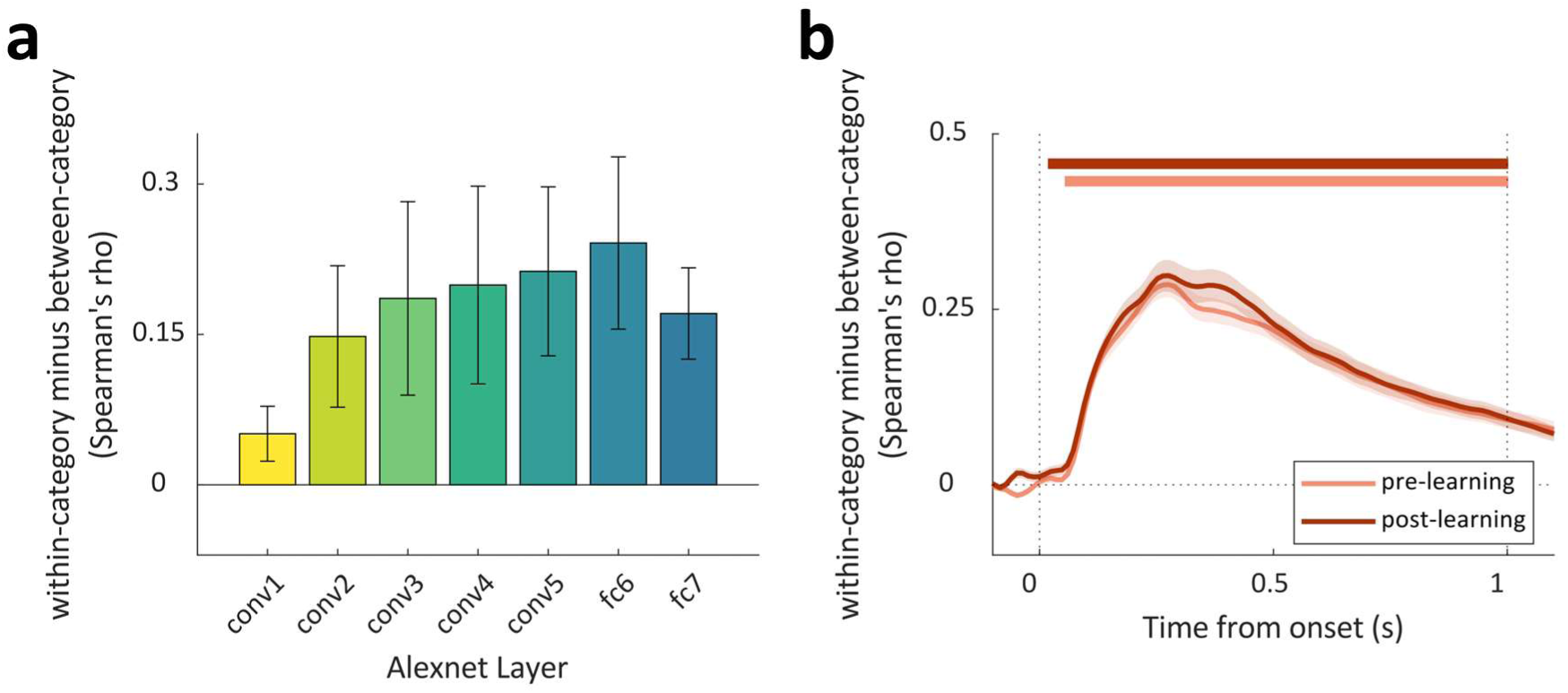
Category representation in DNN and EEG. **(a)** DNN category-specific information (within-category similarity minus between-category similarity) across hidden layers. Error bars indicate standard error across categories (n = 5); **(b)** Neural category-specific information in the perceptual task. Shaded areas indicate standard error across participants. Horizontal lines indicate significant clusters in time (*p*_cluster_ < 0.001, N = 26, two-tailed cluster permutation tests).

**Extended Data Figure 5.**
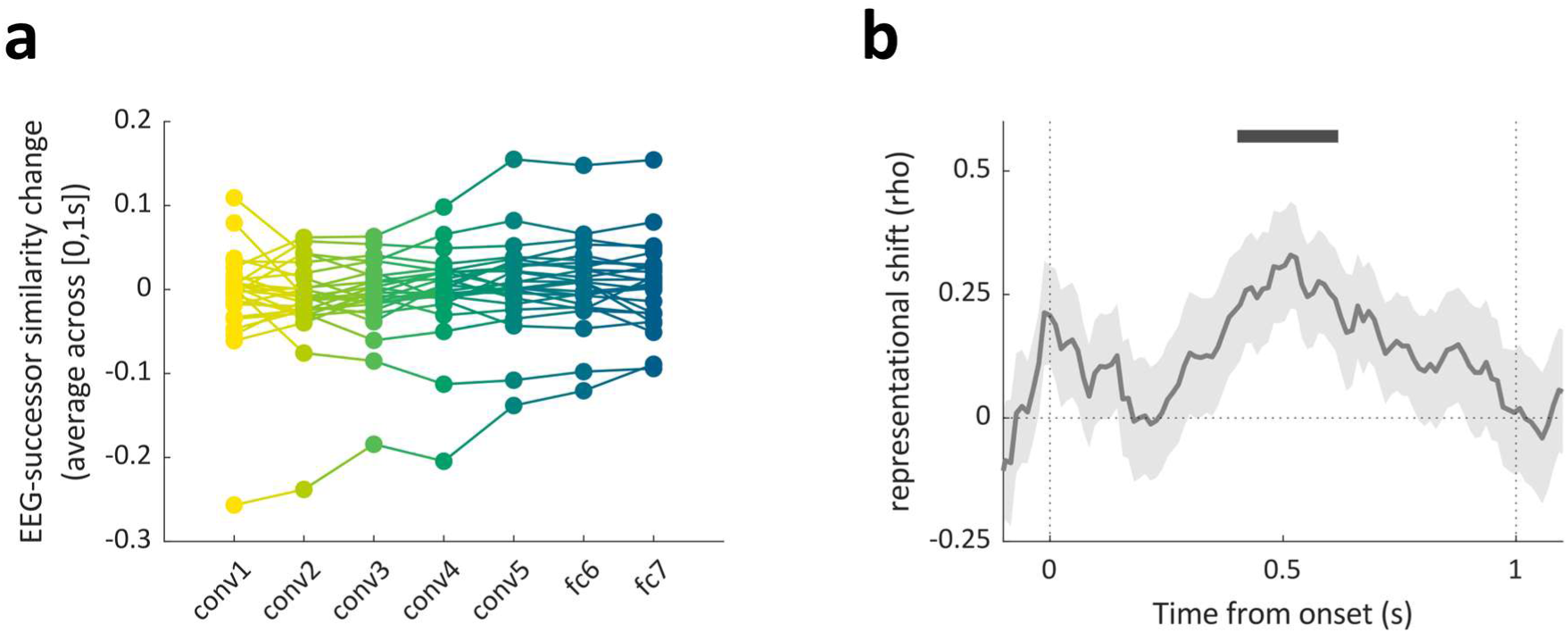
Representational changes between EEG and DNN for successor images. **(a)** Individual participant changes in EEG-successor similarity across DNN layers, averaged across the 0-1s time window. Each line represents one participant (from yellow to blue: conv1, conv2, conv3, conv4, conv5, fc6, fc7); **(b)** Time-resolved correlation between layer hierarchy and representational changes. Shading indicates standard error across participants. Correlations were Fisher z-transformed for statistical testing, with extreme values (>0.99 or <-0.99) winsorized. Horizontal line indicates significant clusters in time (*p*_cluster_ = 0.018, N =26, two-tailed cluster permutation tests).

**Extended Data Figure 6.**
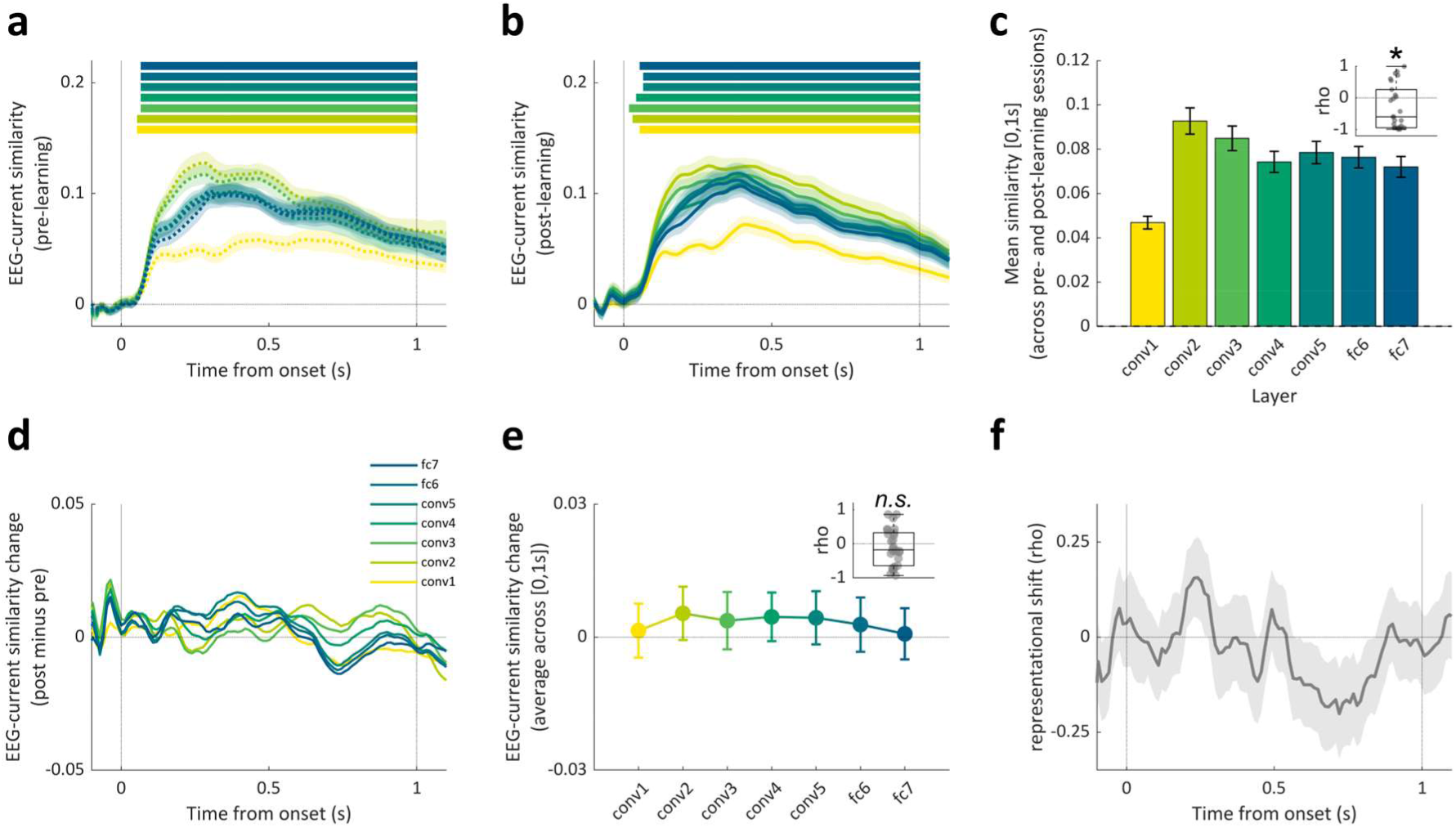
Representational similarity analyses between EEG and DNN for currently displayed images. **(a)** Spearman’s correlations (EEG-current similarity) between EEG similarity matrix in the pre-learning session and DNN current similarity matrices of each layer (from yellow to blue: conv1, conv2, conv3, conv4, conv5, fc6, fc7). Horizontal lines indicate significant correlations (*p*_cluster_ < 0.001, N = 26, two-tailed cluster permutation tests); **(b)** EEG-current similarity in the post-learning session (*p*_cluster_ < 0.001, N = 26, two-tailed cluster permutation tests); **(c)** Average EEG-current similarity across the two sessions. Bar plot shows mean similarity for each DNN layer in the 0-1s time window. Error bars indicate standard error across participants. Inset plot shows the correlations between average similarity and layer hierarchy (from conv2 to fc7). Correlations were Fisher z-transformed for statistical testing, with extreme values (>0.99 or <-0.99) winsorized. * indicates significant difference from zero (*p* = 0.033, two-tailed t-test); **(d)** EEG-current similarity change from pre- to post-learning perceptual task across time. No significant difference was found between the two sessions (*p*_cluster_ > 0.800, N = 26, two-tailed cluster permutation tests); **(e)** Average EEG-current similarity change within the 1-second image presentation time window. Inset plot shows the correlations between average similarity change and layer hierarchy (from conv1 to fc7). *n.s.* indicates non-significant (*p* = 0.357, two-tailed t-test); **(f)** Time-resolved correlation between layer hierarchy and similarity changes (representational shift). Shaded areas indicate standard error across participants. No significant difference was found between correlation values and zero (*p*_cluster_ > 0.800, N = 26, two-tailed cluster permutation tests).

**Extended Data Figure 7.**
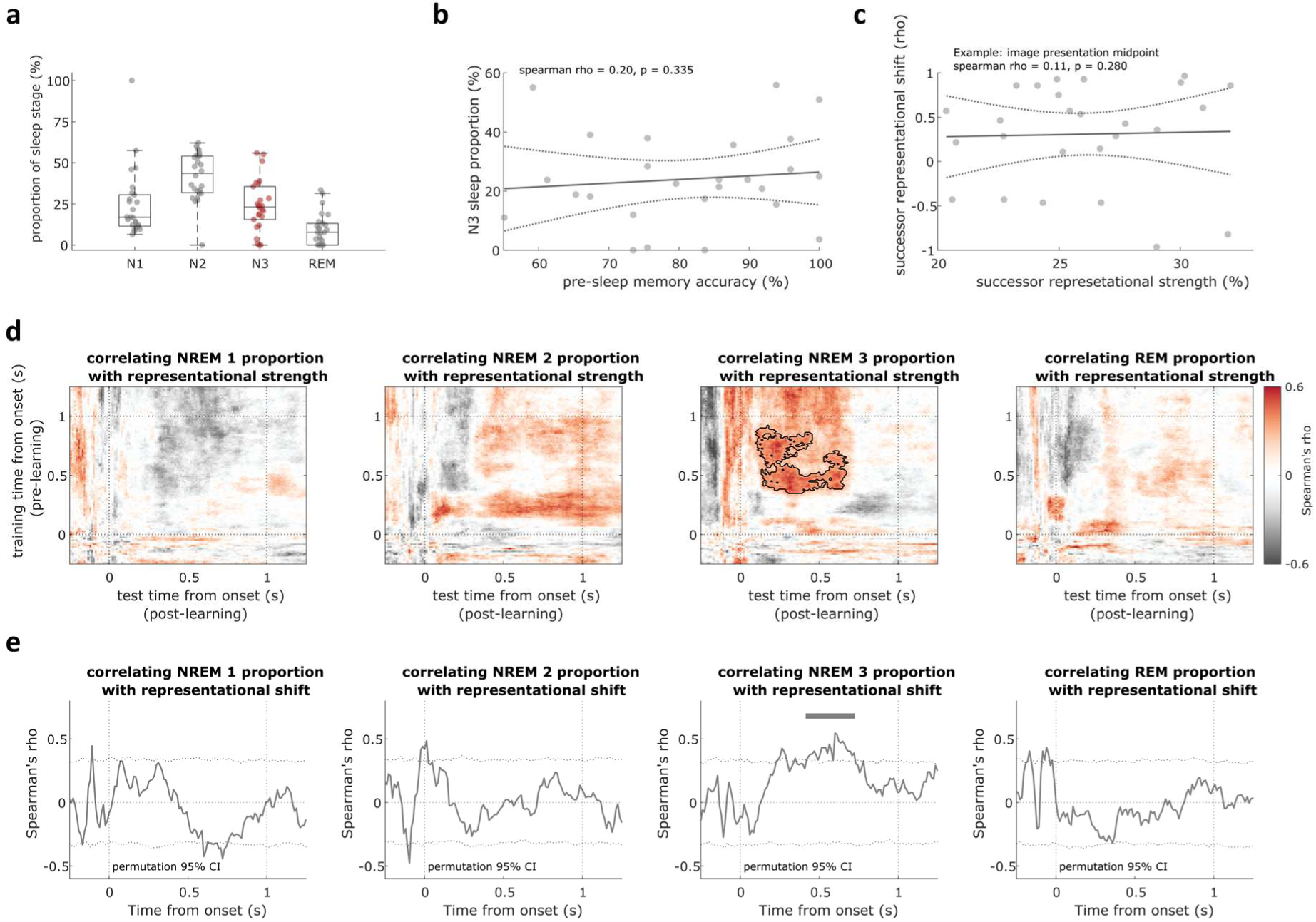
Correlations between successor representational changes and the proportions of sleep stages. **(a)** The proportions of all sleep stages during an approximately 2-hour nap across participants. Each dot represents a participant; **(b)** Correlation between pre-sleep memory accuracy and slow-wave sleep proportion (NREM3; **(c)** Correlation between successor representational strength and shift. The solid line in panels b and c shows the best-fit linear regression line. The dashed lines indicate the 95% confidence bounds for the fitted line; **(d)** Correlation between the proportions of sleep stages and successor representational strength (from left to right: NREM1, NREM2, NREM3, and REM). Black clusters indicate significant correlation (NREM3: *p*_cluster_ = 0.047; Other stages: *p*_cluster_ > 0.145; N = 26, one-tailed cluster permutation tests); **(e)** Correlation between the proportions of sleep stages and successor representational shift (from left to right: NREM1, NREM2, NREM3, and REM). Dashed line indicates 95% CI of 1,000 bootstrapping samples. Horizontal lines indicate significant correlation (NREM3: *p*_cluster_ = 0.013; Other stages: *p*_cluster_ > 0.341; N = 26, one-tailed cluster permutation tests).

## References

Büchel, P. K., Klingspohr, J., Kehl, M. S., & Staresina, B. P. (2024). Brain and eye movement dynamics track the transition from learning to memory-guided action. Current Biology, 34(21), 5054–5061.e4. 10.1016/j.cub.2024.09.063

Buzsáki, G. (2015). Hippocampal sharp wave-ripple: A cognitive biomarker for episodic memory and planning. Hippocampus, 25(10), 1073–1188. 10.1002/hipo.22488

Buzsáki, G., & Tingley, D. (2018). Space and Time: The Hippocampus as a Sequence Generator. Trends in Cognitive Sciences, 22(10), 853–869. 10.1016/j.tics.2018.07.006

Cichy, R. M., Khosla, A., Pantazis, D., Torralba, A., & Oliva, A. (2016). Comparison of deep neural networks to spatio-temporal cortical dynamics of human visual object recognition reveals hierarchical correspondence. Scientific Reports, 6(1), 27755. 10.1038/srep27755

Cowan, E. T., Schapiro, A. C., Dunsmoor, J. E., & Murty, V. P. (2021). Memory consolidation as an adaptive process. Psychonomic Bulletin & Review, 28(6), 1796–1810. 10.3758/s13423-021-01978-x

Dayan, P. (1993). Improving generalization for temporal difference learning: The successor representation. Neural Computation, 5(4), 613–624. 10.1162/neco.1993.5.4.613

Diamond, N. B., Simpson, S., Baena, D., Murray, B., Fogel, S., & Levine, B. (2025). Sleep selectively and durably enhances memory for the sequence of real-world experiences. Nature Human Behaviour, 1–12. 10.1038/s41562-025-02117-5

Diekelmann, S., & Born, J. (2010). The memory function of sleep. Nature Reviews Neuroscience, 11(2), 114–126. 10.1038/nrn2762

Gershman, S. J. (2018). The Successor Representation: Its Computational Logic and Neural Substrates. Journal of Neuroscience, 38(33), 7193–7200. 10.1523/JNEUROSCI.0151-18.2018

Gershman, S. J., Moore, C. D., Todd, M. T., Norman, K. A., & Sederberg, P. B. (2012). The successor representation and temporal context. Neural Computation, 24(6), 1553–1568. 10.1162/NECO_a_00282

Gilboa, A., & Moscovitch, M. (2021). No consolidation without representation: Correspondence between neural and psychological representations in recent and remote memory. Neuron, 109(14), 2239–2255. 10.1016/j.neuron.2021.04.025

Güçlü, U., & van Gerven, M. A. J. (2015). Deep Neural Networks Reveal a Gradient in the Complexity of Neural Representations across the Ventral Stream. The Journal of Neuroscience: The Official Journal of the Society for Neuroscience, 35(27), 10005–10014. 10.1523/JNEUROSCI.5023-14.2015

Goldstein, A., Zada, Z., Buchnik, E., Schain, M., Price, A., Aubrey, B., Nastase, S. A., Feder, A., Emanuel, D., Cohen, A., Jansen, A., Gazula, H., Choe, G., Rao, A., Kim, C., Casto, C., Fanda, L., Doyle, W., Friedman, D., … Hasson, U. (2022). Shared computational principles for language processing in humans and deep language models. Nature Neuroscience, 25(3), 369–380. 10.1038/s41593-022-01026-4

Greco, A., Moser, J., Preissl, H., & Siegel, M. (2024). Predictive learning shapes the representational geometry of the human brain. Nature Communications, 15(1), 9670. 10.1038/s41467-024-54032-4

Hindy, N. C., Ng, F. Y., & Turk-Browne, N. B. (2016). Linking pattern completion in the hippocampus to predictive coding in visual cortex. Nature Neuroscience, 19(5), 665–667. 10.1038/nn.4284

Howard, M. W., & Kahana, M. J. (2002). A Distributed Representation of Temporal Context. Journal of Mathematical Psychology, 46(3), 269–299. 10.1006/jmps.2001.1388

John, T., Zhou, Y., Aljishi, A., Rieck, B., Turk-Browne, N. B., & Damisah, E. C. (2025). Representation of visual sequences in the tuning and topology of neuronal activity in the human hippocampus (p. 2025.03.04.641300). bioRxiv. 10.1101/2025.03.04.641300

Kriegeskorte, N., Mur, M., & Bandettini, P. A. (2008). Representational similarity analysis— Connecting the branches of systems neuroscience. Frontiers in Systems Neuroscience, 2. 10.3389/neuro.06.004.2008

Krizhevsky, A., Sutskever, I., & Hinton, G. E. (2012). ImageNet Classification with Deep Convolutional Neural Networks. In F. Pereira, C. J. Burges, L. Bottou, & K. Q. Weinberger (Eds.), Advances in Neural Information Processing Systems (Vol. 25).

Kumaran, D., Hassabis, D., & McClelland, J. L. (2016). What Learning Systems do Intelligent Agents Need? Complementary Learning Systems Theory Updated. Trends in Cognitive Sciences, 20(7), 512–534. 10.1016/j.tics.2016.05.004

Latchoumane, C.-F. V., Ngo, H.-V. V., Born, J., & Shin, H.-S. (2017). Thalamic Spindles Promote Memory Formation during Sleep through Triple Phase-Locking of Cortical, Thalamic, and Hippocampal Rhythms. Neuron, 95(2), 424–435.e6. 10.1016/j.neuron.2017.06.025

Liu, J., Zhang, H., Yu, T., Ni, D., Ren, L., Yang, Q., Lu, B., Wang, D., Heinen, R., Axmacher, N., & Xue, G. (2020). Stable maintenance of multiple representational formats in human visual short-term memory. Proceedings of the National Academy of Sciences, 117(51), 32329–32339. 10.1073/pnas.2006752117

Liu, J., Zhang, H., Yu, T., Ren, L., Ni, D., Yang, Q., Lu, B., Zhang, L., Axmacher, N., & Xue, G. (2021). Transformative neural representations support long-term episodic memory. Science Advances, 7(41), eabg9715. 10.1126/sciadv.abg9715

Mölle, M., Marshall, L., Gais, S., & Born, J. (2004). Learning increases human electroencephalographic coherence during subsequent slow sleep oscillations. Proceedings of the National Academy of Sciences of the United States of America, 101(38), 13963–13968. 10.1073/pnas.0402820101

Momennejad, I., Russek, E. M., Cheong, J. H., Botvinick, M. M., Daw, N. D., & Gershman, S. J. (2017). The successor representation in human reinforcement learning. Nature Human Behaviour, 1(9), 680–692. 10.1038/s41562-017-0180-8

Nieh, E. H., Schottdorf, M., Freeman, N. W., Low, R. J., Lewallen, S., Koay, S. A., Pinto, L., Gauthier, J. L., Brody, C. D., & Tank, D. W. (2021). Geometry of abstract learned knowledge in the hippocampus. Nature, 595(7865), 80–84. 10.1038/s41586-021-03652-7

Niknazar, H., Malerba, P., & Mednick, S. C. (2022). Slow oscillations promote long-range effective communication: The key for memory consolidation in a broken-down network. Proceedings of the National Academy of Sciences of the United States of America, 119(26), e2122515119. 10.1073/pnas.2122515119

Oostenveld, R., Fries, P., Maris, E., & Schoffelen, J.-M. (2011). FieldTrip: Open source software for advanced analysis of MEG, EEG, and invasive electrophysiological data. Computational Intelligence and Neuroscience, 2011, 156869. 10.1155/2011/156869

Petzka, M., Charest, I., Balanos, G. M., & Staresina, B. P. (2021). Does sleep-dependent consolidation favour weak memories? Cortex, 134, 65–75. 10.1016/j.cortex.2020.10.005

Radford, A., Wu, J., Child, R., Luan, D., Amodei, D., & Sutskever, I. (2019). Language Models are Unsupervised Multitask Learners. OpenAI.

Rau, E. M. B., Fellner, M.-C., Heinen, R., Zhang, H., Yin, Q., Vahidi, P., Kobelt, M., Asano, E., Kim-McManus, O., Sattar, S., Lin, J. J., Auguste, K. I., Chang, E. F., King-Stephens, D., Weber, P. B., Laxer, K. D., Knight, R. T., Johnson, E. L., Ofen, N., & Axmacher, N. (2025). Reinstatement and transformation of memory traces for recognition. Science Advances, 11(8), eadp9336. 10.1126/sciadv.adp9336

Reddy, L., Poncet, M., Self, M. W., Peters, J. C., Douw, L., van Dellen, E., Claus, S., Reijneveld, J. C., Baayen, J. C., & Roelfsema, P. R. (2015). Learning of anticipatory responses in single neurons of the human medial temporal lobe. Nature Communications, 6(1), 8556. 10.1038/ncomms9556

Roüast, N. M., & Schönauer, M. (2023). Continuously changing memories: A framework for proactive and non-linear consolidation. Trends in Neurosciences, 46(1), 8–19. 10.1016/j.tins.2022.10.013

Schapiro, A. C., Kustner, L. V., & Turk-Browne, N. B. (2012). Shaping of Object Representations in the Human Medial Temporal Lobe Based on Temporal Regularities. Current Biology, 22(17), 1622–1627. 10.1016/j.cub.2012.06.056

Schreiner, T., Griffiths, B. J., Kutlu, M., Vollmar, C., Kaufmann, E., Quach, S., Remi, J., Noachtar, S., & Staudigl, T. (2024). Spindle-locked ripples mediate memory reactivation during human NREM sleep. Nature Communications, 15(1), 5249. 10.1038/s41467-024-49572-8

Schreiner, T., Petzka, M., Staudigl, T., & Staresina, B. P. (2021). Endogenous memory reactivation during sleep in humans is clocked by slow oscillation-spindle complexes. Nature Communications, 12(1), 3112. 10.1038/s41467-021-23520-2

Stachenfeld, K. L., Botvinick, M. M., & Gershman, S. J. (2017). The hippocampus as a predictive map. Nature Neuroscience, 20(11), 1643–1653. 10.1038/nn.4650

Staresina, B. P. (2024). Coupled sleep rhythms for memory consolidation. Trends in Cognitive Sciences, 28(4), 339–351. 10.1016/j.tics.2024.02.002

Tacikowski, P., Kalender, G., Ciliberti, D., & Fried, I. (2024). Human hippocampal and entorhinal neurons encode the temporal structure of experience. Nature, 635(8037), 160–167. 10.1038/s41586-024-07973-1

Tarder-Stoll, H., Baldassano, C., & Aly, M. (2024). The brain hierarchically represents the past and future during multistep anticipation. Nature Communications, 15(1), 9094. 10.1038/s41467-024-53293-3

Treder, M. S. (2020). MVPA-Light: A Classification and Regression Toolbox for Multi-Dimensional Data. Frontiers in Neuroscience, 14. 10.3389/fnins.2020.00289

Umbach, G., Kantak, P., Jacobs, J., Kahana, M., Pfeiffer, B. E., Sperling, M., & Lega, B. (2020). Time cells in the human hippocampus and entorhinal cortex support episodic memory. Proceedings of the National Academy of Sciences of the United States of America, 117(45), 28463–28474. 10.1073/pnas.2013250117

Vallat, R., & Walker, M. P. (2021). An open-source, high-performance tool for automated sleep staging. eLife, 10, e70092. 10.7554/eLife.70092

Wilson, M. A., & McNaughton, B. L. (1994). Reactivation of Hippocampal Ensemble Memories During Sleep. Science, 265(5172), 676–679. 10.1126/science.8036517

Yamins, D. L. K., & DiCarlo, J. J. (2016). Using goal-driven deep learning models to understand sensory cortex. Nature Neuroscience, 19(3), 356–365. 10.1038/nn.4244

Zhou, C. Y., Talmi, D., Daw, N. D., & Mattar, M. G. (2025). Episodic retrieval for model-based evaluation in sequential decision tasks. Psychological Review, 132(1), 18–49. 10.1037/rev0000505

